# Fast adaptation in invertebrate looming-sensitive descending neurons

**DOI:** 10.1101/2025.02.10.637552

**Authors:** Katja Sporar Klinge, Karin Nordström

**Affiliations:** South Australia Invertebrate Vision Collective, Flinders Health and Medical Research Institute, Flinders University, GPO Box 2100, Adelaide, SA, 5001, Australia; Department of Medical Cell Biology, Uppsala University, 75123 Uppsala, Sweden

## Abstract

Motion vision plays a crucial role in guiding dynamic behaviors, such as determining when to escape from predators, pursue prey, navigate obstacles, or adjust flight patterns during migration. Adaptation to repetitive motion stimuli is a crucial aspect of this process, allowing animals to efficiently process new stimuli while avoiding sensory overload. This helps animals remain responsive to novel or important stimuli, ensuring appropriate behavioral reactions. Adaptation to looming stimuli, which often signal an approaching threat through the rapid expansion of an object’s image on the retina, allows animals to distinguish harmless from harmful stimuli. While neural adaptation has been extensively studied in the fly’s optic lobes, less is known about how descending neurons, which link the optic lobes to the motor centers in the thoracic ganglia, adapt. To address this gap, we investigate adaptation in looming-sensitive descending neurons in the hoverfly *Eristalis tenax*. Using intracellular recordings, we show that these descending neurons adapt to looming stimuli with inter-stimulus intervals of 1-3 s. We show that the level of adaptation depends on the ISI, with shorter intervals leading to greater adaptation. Specifically, we find that adaptation leads to decreased response duration, with a pronounced delayed response onset. We identified descending neurons that responded to looming stimuli either unilaterally or bilaterally and used this to show that most of the adaptation takes place within the neuron itself, rather than its pre-synaptic inputs. Finally, we found that the wing beat amplitude of tethered hoverflies did not appear to adapt to repetitive looming stimuli.

**Significance Statement:** Rapidly expanding looming objects often signal approaching threats, and appropriate detection is therefore critical for survival across species. Adaptation to repetitive looming cues allows animals to filter out irrelevant information, ensuring they respond only to significant threats. While adaptation has been well studied in the brain, how descending neurons, which connect the brain with the body, adapt to looming stimuli remains poorly understood. Here, we investigate looming-sensitive descending neurons in the hoverfly *Eristalis tenax* and show that they adapt strongly when looming stimuli are repeated in quick succession. We show that most of the adaptation takes place within the descending neuron itself. Our research establishes a foundation for future exploration of the neural mechanisms underlying behavior in constantly changing environments.

## Introduction

Adaptation, which leads to a reduced response to repetitive stimuli, is a fundamental feature of biological systems, enabling maintained sensitivity to novel and behaviorally relevant stimuli, while filtering out redundant or irrelevant signals. Adaptation thus allows neural circuits to prioritize new information while conserving energy by not responding to repetitive stimuli, which are more likely harmless. In visual systems, adaptation is well-documented in e.g. photoreceptors and optic lobe neurons. Photoreceptor adaptation enables responses to changing light intensities and environmental conditions, such as shifting from daylight to dim light (Laughlin, 1989; Van Hateren, 1992), whereas optic lobe neurons adapt to continuous widefield motion, such as the optic flow generated during navigation (Nordström and O’Carroll, 2009; Rankin et al., 2009; Borst et al., 2010).

Looming stimuli are characterized by the rapid expansion of an object’s image, often signaling an approaching threat, making early detection crucial for survival. Across a wide range of species, from insects to vertebrates, looming stimuli trigger a range of behaviors, such as escape, avoidance, freezing, or other defensive reactions (Hemmi and Merkle, 2009; Yilmaz and Meister, 2013; Von Reyn et al., 2014; Peek and Card, 2016; Branco and Redgrave, 2020). For example, crabs adapt to looming stimuli repeated with 1 minute inter-stimulus intervals (ISIs), with a progressive reduction in running distance and increased response latency (Oliva et al., 2007). Zebrafish larvae adapt to repeated dark looming stimuli by reducing their stereotypical escape swims (Mancienne et al., 2021; Marquez-Legorreta et al., 2022). Zebrafish adaptation is mediated by two neural pathways: a looming-specific channel that processes expanding edges to trigger escapes, and a dimming-specific channel that inhibits escape responses under contextual modulation (Fotowat and Engert, 2023).

Neural circuits encoding the behavioral responses to looming stimuli have been identified in a variety of invertebrate and vertebrates (Fotowat and Gabbiani, 2011; Herberholz and Marquart, 2012; Yilmaz and Meister, 2013; Dunn et al., 2016; Peek and Card, 2016; Ache et al., 2019; Branco and Redgrave, 2020). In vertebrates, the optic tectum (or superior colliculus in mammals) is the main visual area for processing looming stimuli (Westby et al., 1990; Branco and Redgrave, 2020). In *Drosophila*, the timing of a single spike in the giant fiber (GF) descending neuron determines whether the fly performs a short or long take-off escape (Von Reyn et al., 2014). In locusts, the optic lobe lobula giant movement detector (LGMD), and its post-synaptic counterpart, the descending contralateral movement detector (DCMD), mediate robust steering maneuvers and escape jumps (Gabbiani et al., 1999; Fotowat and Gabbiani, 2011), while looming sensitive neurons in the central complex affect the direction of the escape (Rosner and Homberg, 2013). Locust DCMD neurons adapt more to repetitive looming stimuli (Matheson et al., 2004; Gray, 2005) when these have an ISI of 4 s compared to 34 s. Furthermore, adapted DCMDs still respond if the looming object approaches along a new trajectory, or if a larger object approaches along the same trajectory (Gray, 2005). This was interpreted as adaptation occuring pre-synaptic to the DCMD, thereby allowing maintained sensitivity to novel stimuli.

While locust DCMDs adapt to repetitive stimuli (Gabbiani et al., 1999; Gray, 2005; Peron et al., 2009; Gray et al., 2010), insects have many different looming-sensitive descending neurons (see e.g. Nicholas et al., 2023), and it is unknown how these adapt. This is important as descending neurons link the sensory centers in the head with the motor command centers in the thoracic ganglia (Hsu and Bhandawat, 2016; Namiki et al., 2018). To address this, we quantified neural and behavioral responses to looming stimuli in hoverflies (*Eristalis tenax*), using different ISIs. We found that looming-sensitive descending neurons adapt strongly when the ISI is 1 or 3 s, but not when it is 10 or 20 s, and that most of this adaptation is intrinsic to the neuron. Interestingly, we found that the wing beat amplitude (WBA) of tethered hoverflies did not adapt to similar stimuli.

## Material and Methods

### Experimental Design

#### Animals

*Eristalis tenax* hoverflies were reared and housed as described previously (Nicholas et al., 2018). In short, pregnant female *Eristalis tenax* hoverflies were collected from the Wittunga Botanic Garden in Adelaide, South Australia, and their larvae raised at room temperature in a rabbit dung slurry. Adult hoverflies were housed in bugdorms (24.5 cm cube) and kept at room temperature with unrestricted access to water, sugar, and pollen. The female hoverflies used for electrophysiology were 28 – 129 days old and for behavior they were 11 – 31 days old.

#### Intracellular electrophysiology

Recordings were made from 66 looming-sensitive descending neurons in 63 female *Eristalis tenax* hoverflies. At experimental time the hoverflies were carefully attached to a plastic holder using a beeswax and resin mixture at room temperature, positioned ventral side up. A small incision was made at the anterior end of the thorax to expose the cervical connective. If the gut exhibited excessive movement or pulsation, it was partially pulled out. The trachea covering the cervical connective were carefully removed using forceps. If the recording area appeared dry, a small amount of PBS (Sigma-Aldrich) was added to the dissected region. A small wire hook was placed under the cervical connective for mechanical support, and a silver wire was inserted into the thoracic cavity as a reference electrode.

Aluminosilicate electrodes were pulled using a Sutter P-1000 micropipette puller (Sutter instruments, San Francisco), resulting in resistances of 40 to 70 MΩ. We either filled the electrodes with 1 M KCl or immersed them, tip pointing upwards, into an Eppendorf tube containing a 3% neurobiotin solution (3 mg of neurobiotin in 100 μL of 1 M KCl) for several minutes, or until the electrode tip was filled. Following this, the electrodes were backfilled with 1 M KCl using a syringe, leaving an air bubble between the filled tip and the KCl solution. The electrodes were placed in an acrylic electrode holder and using a PM-10 Piezo translator (World Precision Instruments, Inc), they were carefully inserted into the cervical connective.

The data were amplified using an Axoclamp-2B amplifier (Axon Instruments, Australia), and 50 Hz noise was minimized with a Humbug (Quest Scientific, Canada). Data acquisition and digitization were performed at 10 kHz using an NI USB-6210 16-bit data acquisition card (National Instruments) and the MATLAB Data Acquisition Toolbox.

#### Tethered flight

Female hoverflies were tethered to a needle (BD Microlance 23G x 1 1/4” - 0.6 x 30 mm Blue hypodermic needles) at a 32° angle using a mixture of bee’s wax and resin. The needle was connected to a syringe (BD tuberculin syringe, 1 ml). To encourage flying we provided airflow manually (Ogawa et al., 2025) and once the hoverfly was flying consistently (10 out of 28 hoverflies), we placed it in the center of the arena. Only flies that maintained continuous flight throughout each presentation of repetitive looming stimuli (see below) were included in the analysis.

The hoverflies were filmed from above at 100 Hz, using a PS3 camera (as in Kumar Kaushik et al., 2020; Ogawa et al., 2025) with a spatial resolution of 240 x 320 pixels. The camera was equipped with an infrared pass filter, and the hoverfly illuminated with infrared lights, and a musou black surface was placed below the hoverfly to maximize the contrast.

#### Visual stimuli

For electrophysiology the fly was positioned 13 cm away from the center of the ViewSonic monitor (ViewSonic, California, USA), with a refresh rate of 165 Hz and a spatial resolution of 2560 x 1440 pixels (width x height), covering 133° of the fly’s visual field in azimuth and 107° in elevation.

For behavior, the hoverfly was placed at 10 cm distance from the center of the linearized monitor (Asus, Taipei, Taiwan). The monitor subtended 1440 x 2560 pixels (width x height), covering 121° of the fly’s visual field in azimuth and 143° in elevation, with a refresh rate of 165 Hz.

We used custom written scripts based on the Psychophysics toolbox (Brainard, 1997; Pelli, 1997) in Matlab (Mathworks) to generate the following visual stimuli:

#### Looming stimulus and motion-free control

We used previously described methods to generate looming stimuli (Fotowat and Gabbiani, 2007; Nicholas et al., 2020) using an *l/|v|* of 8 ms for electrophysiology. This resulted in a stimulus where a small black circle (1 °) appeared on a white monitor and expanded to a final diameter (103°) over 1 s. It remained in full size on the monitor for 1 s. A motion-free control was a black circle with diameter of 103° appearing and remaining on the screen for 2 s.

#### Repetitive looming stimuli

For electrophysiology, above-described looming stimulus was shown in a sequence of at least 7 repetitions. Thus, a circular black disc expanded over 1 s from 1° to 103°, remained on the screen for 1 s, and then disappeared for a given ISI. The ISIs were 1, 3, 10 or 20 s. In between the sets of repetitions there was a minimum gap of at least 20 s during which different stimuli were shown, such as wide field grating.

For behavior, a circular black disc expanded over 1 s from 1° to 117°, with an l/|v| of 6 ms. It remained stationary at 117° for 1 s, and then disappeared for an ISI of 1, 3, 5 or 10 s. To keep the hoverflies flying, a starfield stimulus sideslipping to the right at 50 cm/s (Nicholas et al., 2020) was presented for a minimum 20 s between each adaptation series, where the different ISIs were presented in random order.

#### Local adaptation to looming

We presented two looming stimuli, one on the left and the other on the right side of the monitor. The looming stimuli were presented 9 times, alternating between the sides. The first stimulation was always on the neuron’s preferred side. All neurons were then manually classified as having a preferred (P) or non-preferred (N) side, based on the responses to the mapping of the looming receptive field. The looming stimuli had an *l/|v|* of 8 ms, growing in size 1° to 103° over 1 s, remaining in full size on the monitor for 1 s, with an ISI of 1 s.

#### Looming receptive field

To map the looming receptive field, the visual monitor was divided into a 4×4 grid. Small looming stimuli (*l/|v|* = 8 ms, 1° to 40° expansion over 1 s, remaining on the screen for 1 s, ISI of 1 s) were presented at randomized locations within the grid.

### Statistical analysis

#### Data analysis, electrophysiology

When a neuron was impaled, a drop in voltage was observed. Only neurons with a stable resting potential and spikes of at least 10 mV amplitude were used. To generate spike histograms, we applied a 50 ms square-wave filter with a resolution of 0.1 ms. The maximum (Fig. 1) or mean (Fig. 2-4) response for each neuron was then identified. For the mean, we used a time window of 160 ms before and 110 ms after the maximum loom size was reached. Looming-sensitive neurons were identified based on a stronger response to the looming stimulus compared with the motion-free control (Nicholas et al., 2020), by calculating looming selectivity (Fig. 1): *(Resp_MaxLoom_ − Resp_MaxControl_) / (Resp_MaxLoom_ + Resp_MaxControl_)*. We included all neurons that had a looming selectivity above 0.25 (see Nicholas et al., 2020).

**Fig. 1.**
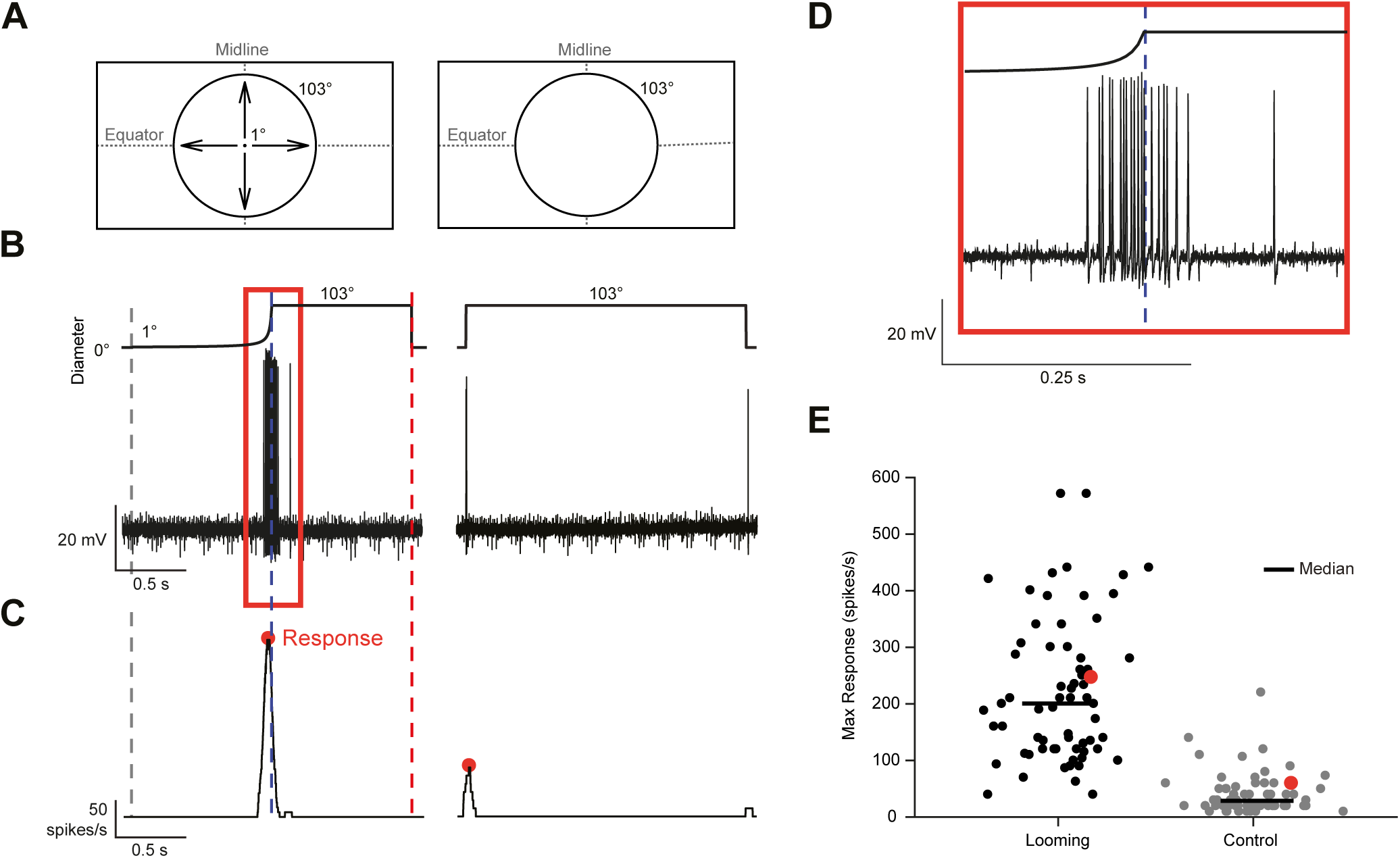
Looming-sensitive descending neurons respond stronger to looming stimuli than to stationary controls. **A)** Pictograms of the looming stimulus (left), with an l/|v| of 8 ms and a final size of 103°, and the stationary control (right), with a diameter of 103°, as projected on the visual monitor. **B)** Raw intracellular recording traces show responses to the looming stimulus (left) and the stationary control (right), with time aligned stimuli shown above. **C)** Smoothed spike histograms (50 ms square-wave filter, 0.1 ms resolution) of the responses in panel B. The red circles indicate the maximum responses. **D)** Magnification of the highlighted response shown in panel B. **E)** Maximum responses across looming-sensitive descending neurons to looming (black) and control (gray) stimuli, with the responses shown in panel C highlighted in red. *N = 63 flies (66 neurons)*.

**Fig. 2.**
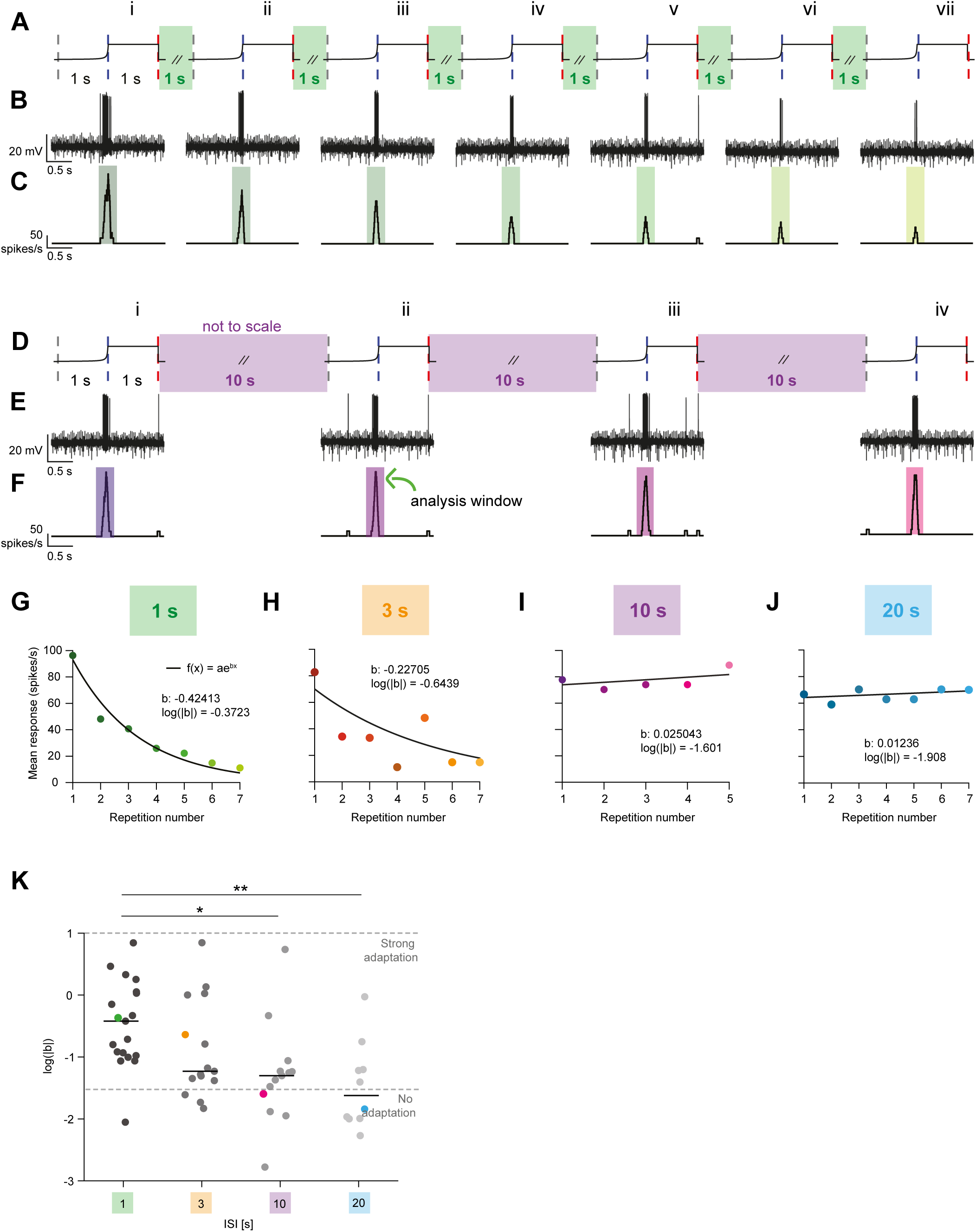
Looming-sensitive descending neurons adapt to repetitive looming stimuli with short inter-stimulus intervals. **A)** Stimulus schematic of repetitive looming stimuli with an inter-stimulus interval (ISI) of 1 s. The looming stimulus had an l/|v| of 8 ms and a final size of 103° (see Fig. 1A, B). **B)** Example intracellular responses corresponding to the schematic in panel A. **C)** Smoothed spike histograms (50 ms square-wave filter, with 0.1 ms resolution), with the analysis windows highlighted in green. **D)** Stimulus schematic of repetitive looming stimuli with 10 s ISI. **E)** Example response traces corresponding to the schematic in panel D. **F)** Smoothed spike histograms, with the analysis windows highlighted in purple. **G-J)** Mean responses to repetitive looming stimuli with ISI of 1 s (G), 3 s (H), 10 s (I), or 20 s (J) from the same neuron as in panels A-F. The mean responses are plotted as a function of repetition number and fitted with an exponential function, *f(x) = ae^bx^*. **K)** The time course of adaptation, defined as the logarithm of the absolute b values from the exponential fits. Colored dots represent the data shown in panels G, H, I, and J, respectively. Kruskal-Wallis test, p = 0.0014; followed by Dunn’s test with Bonferroni correction: p = 0.3905 for 1 – 3 s comparison, p = 0.0126 for 1 – 10 s, p = 0.0041 for 1 to 20 s, p = 1.0000 for 3 – 10 s, p = 0.5436 for 3 – 20 s, p = 1.0000 for 10 – 20 s, *N = 10-18*.

Mean responses to the repetitive looming stimuli were fitted with an exponential function (*f(x) = ae^bx^*). We assigned weights to the response from each repetition, giving higher importance to the first repetition by setting its weight to 2, while keeping the weights for all other repetitions to 1. This emphasized the initial response during the exponential fitting process. We focused on the *b*-value from the exponential fit because it represents the change of the response over time. We took the logarithm of the positive *b*-values, *log(|b|)*, a transformation that stabilizes the variance and makes it easier to compare across different datasets.

To investigate the time course of adaptation, we calculated the delta time (*ΔT*), defined as the time difference between the first spike in the last repetition (*t_L1_*) and the first spike in the first repetition (*t_F1_*; *ΔT=t_L1_−t_F1_*, see Fig. 3B, D). To calculate the response duration ratio, we used the difference between the time of the last spike and the time of the first spike of the last repetition (*t_Lend_* and *t_L1_*), divided by the difference between the last and the first spike of the first repetition (*t_Fend_* and *t_F1_*, see Fig. 3C, E), using: *Dur Ratio =* (*t_Lend_−t_L1_)/(t_Fend_−t_F1_)×100*.

**Fig. 3.**
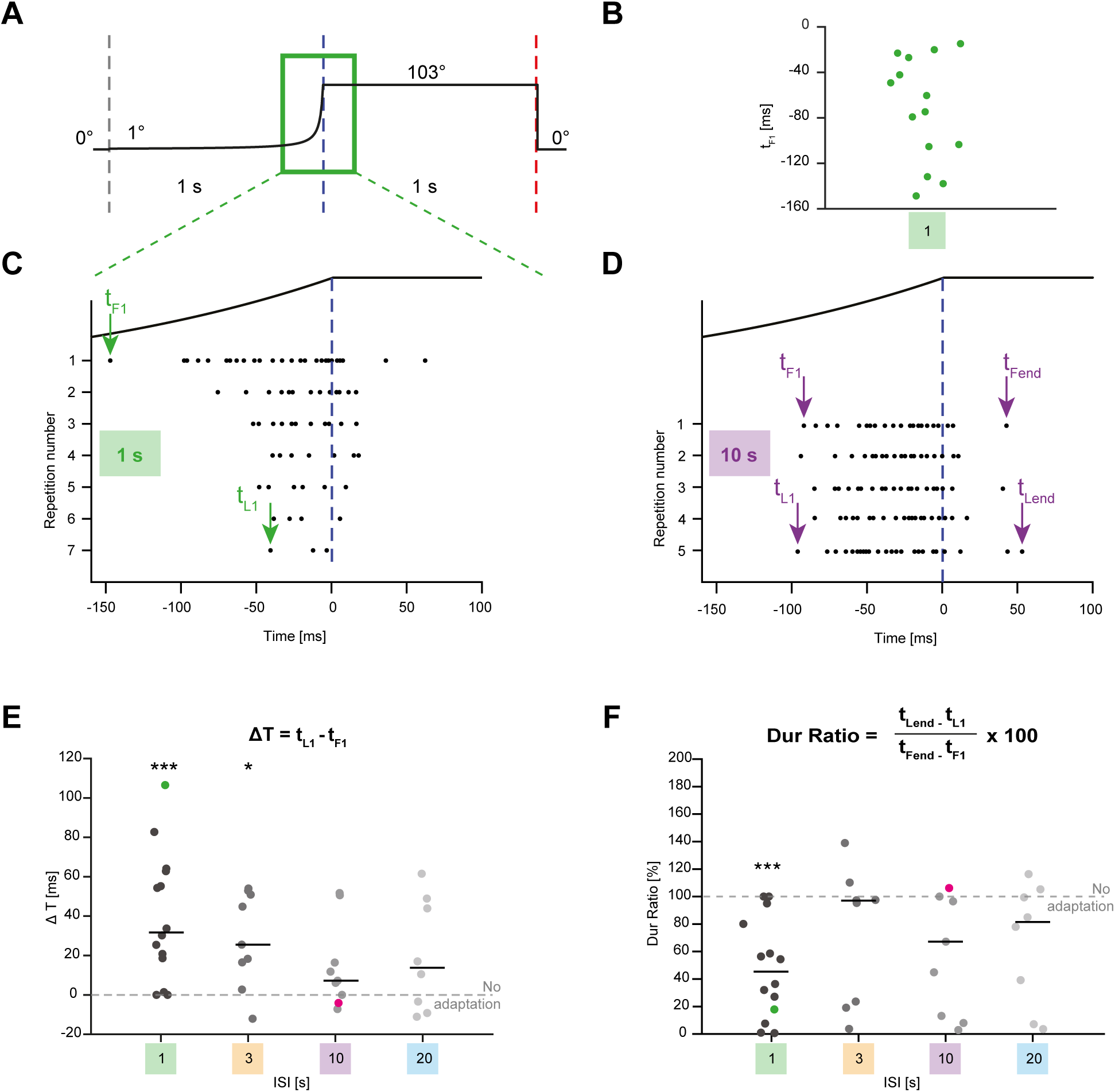
The adaptation of looming-sensitive descending neurons primarily takes place at the start of the response. **A)** Stimulus pictogram with the green box indicating the zoomed-in time window, corresponding to the response raster plots shown in panels C and D. **B)** The time of the first spike in response to the first loom presentation, *t_F1_*, for ISI 1 s. **C)** Example raster plot of a single neuron showing spike times in response to repetitive looming stimuli, with an ISI of 1 s. Each row shows the response to a new repetition. The time of the first spike in response to the first stimulus presentation is denoted as *t_F1_* and the time of the first spike in the last repetition as *t_L1_*. **D)** Example raster plot from the same single neuron showing spike times in response to looming stimuli, with an ISI of 10 s. The time of the last spike in the first repetition as *t_Fend_,* and the time of the last spike in the last repetition as *t_Lend_.* **E)** Delta time (*Δt*) was defined as the difference between the time of the first spike in the last repetition (*t_L1_*) and the time of the first spike in the first loom repetition (*t_F1_*). The dashed gray line indicates no adaptation (*Δt* = 0). For ISI values of 1 s and 3 s, *Δt* is significantly different from 0, suggesting adaptation (one-sample Wilcoxon Signed-Rank test; p = 0.0005, 0.0117, 0.0547, 0.1484). **F)** The duration ratio was defined as *(t_Lend_−t_L1_)/(t_Fend_−t_F1_)×100*. The gray dashed line indicates no adaptation (Dur Ratio = 100 %). When the ISI is 1 s, the duration ratio is significantly different from 100%, indicating adaptation (one-sample Wilcoxon Signed-Rank test; p = 0.0005, 0.6523, 0.3125, 0.1094). In panels E and F, *N = 8-14*.

The hoverfly was placed ventral side up in electrophysiology experiments, but receptive fields are displayed with the dorsal side up (Fig. 4F, G). We plotted the responses to the individual looming stimuli as histograms (Fig. 4F, G), with the color coding indicating the maximum responses at each location. Neurons whose receptive fields covered only one side of the monitor were classified as unilateral, while those with receptive fields extending across both sides of the monitor were manually categorized as bilateral neurons, based on response visualization and spike raster plots in MATLAB. All neurons were then manually classified as having a preferred (P) or non-preferred (N) side.

**Fig. 4.**
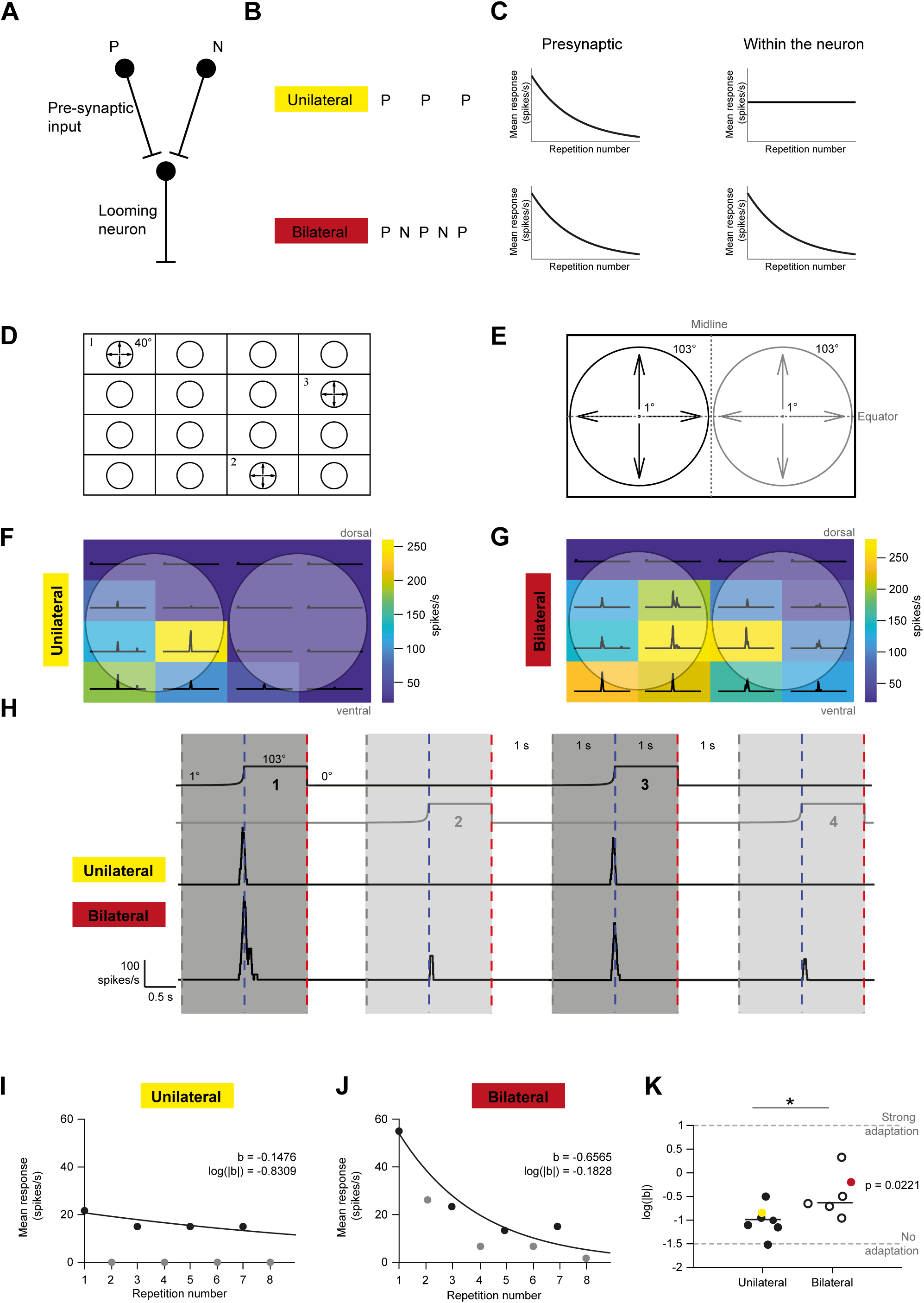
Looming-sensitive descending neurons adapt within the neuron itself. **A)** Adaptation may primarily occur in the pre-synaptic units or within the descending neurons themselves. This can be tested using neurons that have a preferred (P) and a non-preferred (N) side. **B)** Unilateral neurons respond only to their preferred (P) side, whereas bilateral neurons respond to both their preferred (P) and non-preferred side (N). When looming stimuli are presented alternately on the preferred (P) or non-preferred (N) side, unilateral neurons are only excited by preferred side (P) stimulation, whereas bilateral neurons are excited by both sides. **C)** Left: If adaptation occurs entirely in the pre-synaptic units, the response to stimuli presented on the preferred side (P) should be identical whether recorded from unilateral or bilateral neurons. Right: However, if adaptation occurs within the descending neuron itself, the response to stimuli on the preferred side should show less adaptation in unilateral neurons compared to bilateral neurons. **D)** Stimulus design for mapping the looming receptive field. The visual monitor was divided into a 4×4 grid, with small looming stimuli (l/|v| = 8 ms, 40° maximum size, timing as in Fig. 1) presented at randomized locations within the grid. **E)** Stimulus pictogram illustrating the placement of looming stimuli, presented either on the preferred (black, here left) or non-preferred (gray, here right) side of the monitor. **F)** Example unilateral receptive field. The matrix squares are color-coded to represent the maximum response at each location, while the lines within each square depict the spike histogram for that location. Circles indicate the stimulus location at maximum size. **G)** Example bilateral receptive field, showing the strongest responses in the ventral visual field. **H)** Top: Stimulus pictogram showing alternating loom presentations on the preferred (black, repetition 1 and 3) and non-preferred (gray, repetition 2 and 4) sides. Middle: The unilateral neuron responded exclusively to the preferred side stimuli (same neuron as in panel F). Bottom: The bilateral neuron responded to both sides (same neuron as in panel G). **I-J)** Mean responses of the unilateral (I) and bilateral neuron (J) plotted against repetition number. Black data correspond to preferred side stimulation, and gray data to non-preferred side. The exponential function was fitted to the preferred side responses. **K)** Logarithm of absolute *b* values for unilateral and bilateral neurons. Colored points correspond to the neurons shown in panels F, G, H, I, and J. Mann-Whitney U test (p = 0.0221). *N = 6-7*.

#### Data analysis, behavior

We used DeepLabCut (DLC) version 2.3.3 (Nath et al., 2019; Kane et al., 2020) to train a model to track 6 points on the hoverfly (Ogawa et al., 2025). For this, we manually labelled 16 extracted video frames each from videos of six individual animals (3 males, 3 females). We trained the DLC model to label two points along the animal’s longitudinal axis, and two each along the anterior edge of each wing stroke and then used Matlab to calculate the peak downstroke angle, also referred to as the wing beat amplitude (WBA), of each wing (Maimon et al., 2010; Ogawa et al., 2025). For further analysis we used the sum (ΣWBA) of the left and the right WBA.

#### Statistical analysis

All statistical analysis was performed in Matlab. The sample size, statistical test, and *p*-values are indicated in the main text and each figure legend, where *N* refers to the number of neurons or animals. All data are given as median ± standard deviation, unless otherwise mentioned.

As the data were non-normally distributed, we used Kruskal-Wallis test followed by Dunn’s test for multiple comparisons with Bonferroni correction (Fig. 2K), one-Sample Wilcoxon Signed-Rank test (Fig. 3E, F), or a Mann-Whitney U test (Fig. 4K), for statistical analysis.

Statistical significance is indicated by asterisks, with a single asterisk (*) representing *p <* 0.05, double asterisks (**) representing *p <* 0.01, and triple asterisks (***) representing *p <* 0.001.

## Results

### Looming-sensitive descending neurons respond stronger to looming stimuli than to stationary controls

To characterize adaptation of looming-sensitive neurons, we performed intracellular recordings in the cervical connective while visually stimulating the hoverfly (female *Eristalis tenax*). We presented a black looming stimulus on a white screen, with an *l/|v|* of 8 ms and a final size of 103°, and a motion-free control, with a diameter of 103° (Fig. 1A). Looming neurons are defined by their strong responses to looming stimuli and small or no response to a motion-free control (Fig. 1B). We applied a 50 ms square-wave filter with a resolution of 0.1 ms to generate spike histograms (Fig. 1C) and identified the maximum response within an analysis window (160 ms before to 110 ms after the stimulus reaches its maximum size, red rectangle, Fig. 1B, D). Across 66 neurons the response is 200 ± 126 spikes/s to the looming stimulus, compared with 25 ± 36 spikes/s to the control (Fig. 1E). We show the individual responses for all neurons (Fig. 1E) that passed our threshold (looming selectivity > 0.25 (Nicholas et al., 2020), and included these for further analysis.

### Looming-sensitive descending neurons adapt to repetitive looming stimuli with short inter-stimulus intervals

To characterize the adaptation of looming-sensitive neurons, we used seven repetitive looming stimuli. When using a short, 1 s inter-stimulus interval (ISI), a visual inspection of the data from a single neuron clearly shows a response reduction across repetitions (Fig. 2A-C). In contrast, when the ISI is 10 s, there is no clear evidence of response reduction across repetitions in the same example neuron (Fig. 2D-F).

To investigate this observation, we quantified the mean response to each repetition in each neuron (shaded analysis windows, Fig. 2C, F) and fitted an exponential function (*f(x)=ae^bx^*, Fig. 2G-J). The responses as a function of repetition number revealed drastic adaptation when the ISI was 1 or 3 s (Fig. 2G, H, same single neuron data), whereas there was no strong adaptation when the ISI was 10 or 20 s (Fig. 2I, J). For quantification across neurons, we used the logarithm of the absolute *b*-values and plotted these as a function of the different ISIs (Fig. 2K), where *log(|b|)* close to 0 indicates strong adaptation (see e.g. Fig. 2G), whereas *log(|b|)* around −1.5 indicates weak or no adaptation (see e.g. Fig. 2I). Median *log(|b|)* values were - 0.4262 ± 0.6972 for ISI 1 s, −1.2361 ± 0.7965 for ISI 3s, −1.3076 ± 0.8284 for ISI 10 s and - 1.6279 ± 0.6921 for ISI 20 s. This analysis confirms that across neurons, ISIs of 1 or 3 s generate strong adaptation, whereas ISIs of 10 and 20 s generate weak adaptation (Kruskal-Wallis test, p: 0.0014; followed by Dunn’s test with adjusted p-values using Bonferroni correction for multiple comparisons: p = 0.3905 for 1 – 3 s, p = 0.0126 for 1 – 10 s, p = 0.0041 for 1 to 20 s, p = 1.0000 for 3 – 10 s, p = 0.5436 for 3 – 20 s, p = 1.0000 for 10 – 20 s, *N = 10-18*, Fig. 2K).

### The adaptation of looming-sensitive descending neurons primarily takes place at the start of the response

As seen above (Fig. 1B, C, 2A-F), looming neurons tend to give the strongest response when the looming stimulus reaches its maximum size (see also Gabbiani et al., 1999, 2002; Matheson et al., 2004; Gray, 2005; Oliva et al., 2007; Nicholas et al., 2020, 2023). We quantified the time of the first spike (t_F1_) in response to the first repetition of the looming stimuli with an ISI of 1 s, relative to the time of maximum loom size (time = 0 s). The data confirms that the first spikes occur before the maximum loom size (Fig. 3B), no earlier than −160 ms (Fig. 3B-D).

In the analysis above we quantified the mean spike rate within an analysis window (red box, Fig 1B, D; colored shading, Fig. 2C, F). To investigate the mechanisms of adaptation in more detail we examined its time course. Visual inspection of the data from a single neuron shows that when the ISI is short (1 s), the first spike appears progressively later across repetitions (Fig. 3C). In contrast, when the ISI is longer (10 s), there is no such apparent shift in the timing of the first spike (Fig. 3E). To quantify this observation, we calculated the delta time *(ΔT*). If there is no change in the timing of the first spike, *ΔT* will be 0 ms (dashed line, Fig. 3E). Across neurons, we observed that for ISIs of 1 and 3 s, *ΔT* was significantly different from 0 (median 32 ± 32.4 ms and 25.5 ± 23.9 ms, one-Sample Wilcoxon Signed-Rank test, p = 0.0005 and 0.0117), but not for the two longer ISIs (median 7.2 ± 21.9 ms and 13.8 ± 28.3 ms, p = 0.0547 and 0.1484, see Fig. 3E).

Next, to assess whether the response duration decreased with repetition, we calculated the duration ratio. This was defined as the time between the first and last spike in the last repetition, divided by the difference between the last and the first spike of the first repetition (Fig. 3D, F). In this case, a duration ratio of 100% means that there is no change across repetitions (dashed line, Fig. 3F). We found that the duration ratio was significantly different from 100% when the ISI was 1 s (one-Sample Wilcoxon Signed-Rank test, p = 0.0005, median 45.4 ± 35 %, Fig. 3F). For the longer ISIs the duration ratios were 97.3 ± 202 % (3 s), 67.2 ± 43 % (10 s), and 81.4 ± 44 % (20 s, Fig. 3E; one-Sample Wilcoxon Signed-Rank test, p = 0.6523, p = 0.3125, p = 0.1094, respectively). Taken together, these results suggest that adaptation of looming-sensitive descending neurons primarily occurs by delaying the response onset (Fig. 3C, E).

### Adaption mainly takes place within the descending neuron itself

Next, we aimed to investigate whether the adaptation mainly takes place in pre-synaptic units or within the descending neurons themselves (Fig. 4A). For this purpose, we searched for neurons that responded to only one side (unilateral) or both sides (bilateral, Fig. 4B) of the monitor. For each neuron we defined a preferred (P) side based on a bigger response to looming stimuli, and the opposite side as non-preferred (N), which gave no response in unilateral neurons (top, Fig. 4B). We reasoned that if adaptation is entirely presynaptic, the adaptation to stimuli presented on the preferred side (Fig. 4A, B) should be identical whether we recorded from unilateral or bilateral neurons (left, Fig. 4C). In contrast, if the adaptation takes place within the descending neuron itself, the response to stimuli presented on the preferred side should adapt less in unilateral compared with bilateral neurons (right, Fig. 4C), as they have a longer effective ISI (Fig. 4B).

We mapped the receptive field (Fig. 4D) before displaying the alternating stimuli (Fig. 4E) and classified each neuron as either unilateral (example neuron, Fig. 4F) or bilateral (example, Fig. 4G). We then investigated adaptation to looming stimuli alternating between its preferred (P) or non-preferred (N) side (Fig. 4A, B, E, *l/|v|* = 8 ms, 1 s ISI). The example data confirms that the unilateral neuron only responds when the looming stimulus is on its preferred side (Fig. 4F, H), whereas the bilateral neuron responds to looming stimuli on both sides (Fig. 4G, H). In addition, the response to third presentation (“3”, Fig. 4H) is smaller compared to the first presentation (“1”, Fig. 4H) in the bilateral neuron (bottom trace, Fig. 4H) compared with the unilateral neuron (top trace, Fig. 4H).

We quantified the mean response of each neuron within the same analysis window as before and plotted the preferred side responses as a function of repetition number (black data, Fig. 4I, J). As before, we fit this with an exponential function, *f(x)=ae^bx^,* and extracted *log(|b|)* (Fig. 4I, J). Across neurons we found that unilateral neurons adapt less (median *log|b|* −0.99 ± 0.31; Fig. 4I, K) than bilateral neurons do (median (*log|b|)* −0.63 ± 0.49; Mann-Whitney U test, p = 0.0221; Fig. 4K). This suggests that most of the adaptation takes place within the descending neuron itself (compare Fig. 4C with 4I, J).

### Flying hoverflies do not adapt to repetitive looming stimuli

Above, we have shown that looming-sensitive descending neurons adapt when the ISI is 1 or 3 s, but not when it is longer (Fig. 1-3). To determine if hoverfly behavior adapts, we recorded the wing beat amplitude (WBA) of tethered hoverflies. The stimulus was similar to before (*l/|v|* of 6 ms and a final size of 117°, repeated 10 times, Fig. 5A). For each fly, we calculated the WBA (Fig. 5B) for both the left (Fig. 5C, upper) and right wing (Fig. 5C, lower). We found that the WBA increased shortly after the loom reached its maximum size and then decreased slowly over the next few seconds (Fig. 5D, example data). For quantification across flies we calculated the sum of the left and right WBA (ΣWBA = WBA_L_ + WBA_R_) and plotted this as a function of time (Fig. 5F). This analysis shows no sign of adaptation over time. Instead, we found that the hoverflies WBA increased to a similar magnitude to each looming stimulus repetition, regardless of the ISI duration (Fig. 5F).

**Fig. 5.**
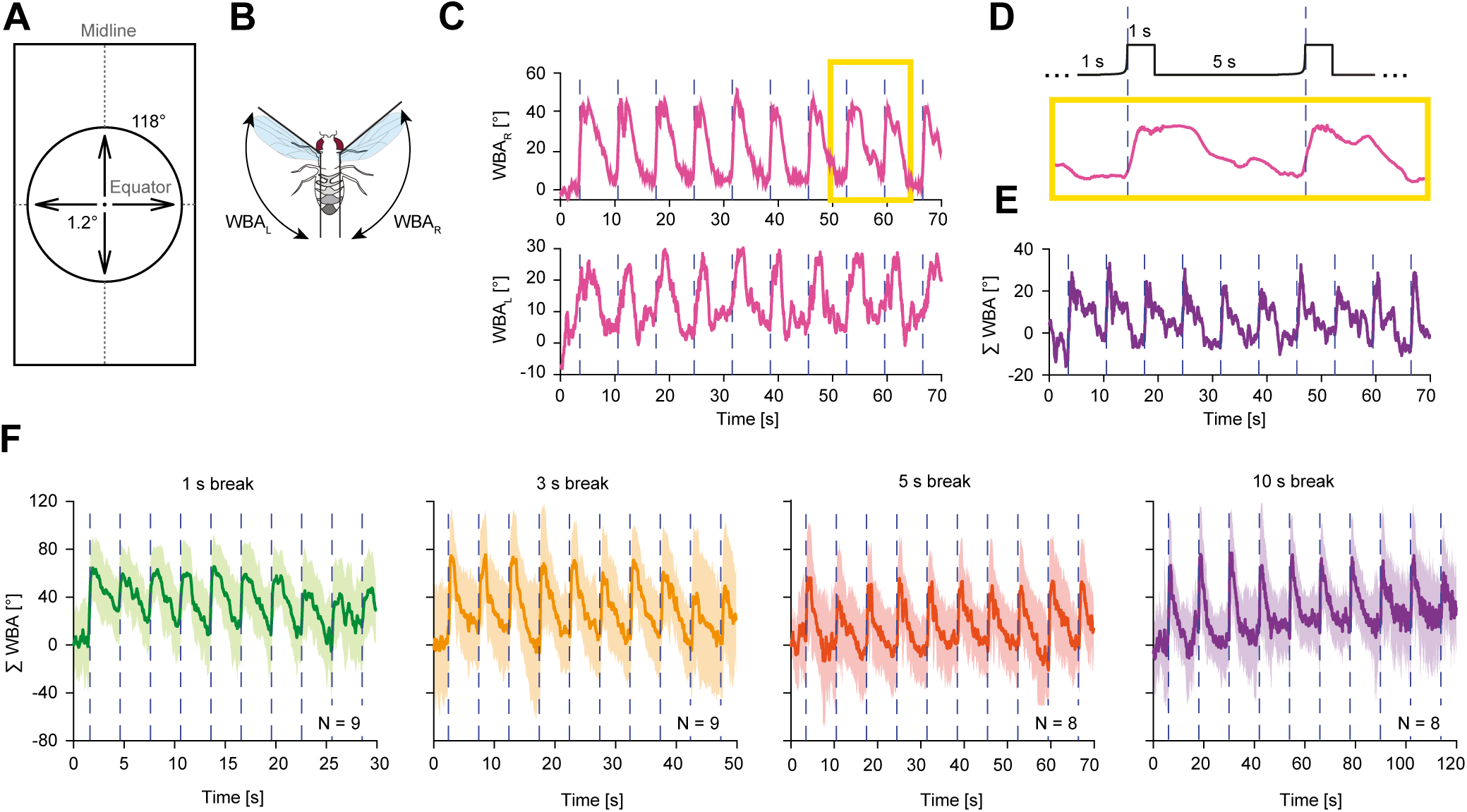
Flying hoverflies do not adapt to repetitive looming stimuli. **A)** Pictogram of a looming stimulus with an l/|v| of 6 ms and a final size of 117°, as projected on the visual monitor in front of the hoverfly. The looming stimulus was repeated ten times with 4 different ISIs. **B)** Pictogram of the wingbeat amplitude (WBA). **C)** Example traces of right (top) and left (bottom) WBA. The traces were normalized to the pre-stimulus WBA. The yellow highlighted area indicates the region magnified in panel D. **D)** Zoomed-in region of the highlighted area in panel C, showing the WBA of the right wing during two looming stimulus repetitions (pictogram at top). **E)** The summed wingbeat amplitude (ΣWBA), defined as WBA_L_ + WBA_R_, of the data shown in panel C. **F)** Average ± standard deviation ΣWBA across flies to looming stimuli presented at ISIs of 1, 3, 5, or 10 s. *N = 8–9*.

## Discussion

In this study, we investigated the responses of looming-sensitive descending neurons in hoverflies (*Eristalis tenax*) to repetitive looming stimulation. Using intracellular recordings, we defined looming neurons based on stronger responses to looming stimuli compared to motion-free controls (Fig. 1). We found that looming-sensitive descending neurons exhibit significant adaptation to repetitive stimulation when inter-stimulus intervals (ISIs) are short (1 or 3 s, Fig. 2). This adaptation is most pronounced at the beginning of the response, with longer latency to the first spike (Fig. 3). Further, we established that adaptation is happening within the descending neuron itself (Fig. 4). Despite neuronal adaptation, behavioral responses remained consistent across all ISIs (Fig. 5). This highlights the challenges in correlating neuronal adaptation with behavior.

### Looming-sensitive descending neurons respond selectively to looming stimuli

As previously, we defined looming-sensitive descending neurons based on their stronger response to looming stimuli than to stationary controls (Fig. 1, and see e.g. Nicholas et al., 2020). This definition works robustly across various species, including insects, birds, and mammals, highlighting a conserved mechanism for detecting potential threats (Peek and Card, 2016). In many insects looming neurons additionally often have the widest diameter among the descending neurons, including the Giant Fiber (GF) in *Drosophila* (Levine and Tracey, 1973; Tanouye and Wyman, 1980; Bacon and Strausfeld, 1986; Mu et al., 2014), and the DCMD in locusts (Burrows and Rowell, 1973; O’shea et al., 1974; Simmons, 1980). The selective response to looming stimuli, together with the large axon diameter, ensures that the behavioral response is fast, and highlights the critical evolutionary role of these neurons in predator detection and collision avoidance (Rind and Simmons, 1999).

Interestingly, however, the looming-sensitive descending neurons in hoverflies appear to constitute a diverse class of neurons (Nicholas et al., 2020), suggesting that they are not direct homologues of the *Drosophila* Giant Fiber (Levine and Tracey, 1973; Tanouye and Wyman, 1980; Bacon and Strausfeld, 1986; Mu et al., 2014) or the locust DMCD (Burrows and Rowell, 1973; O’shea et al., 1974; Simmons, 1980). Indeed, hoverfly looming-sensitive descending neurons have receptive fields with diverse sizes and locations (Nicholas et al., 2020). In addition, we here found a high variability between neurons, both in terms of the response rate (Fig. 1E), but also in their level of adaptation (Fig. 2K, Fig. 3E, F), supporting the suggestion that they come from a wide population, but this needs to be confirmed with morphological fills in the future.

### Adaptation of looming-sensitive descending neurons to repetitive looming stimuli with short inter-stimulus intervals

Our results demonstrated that looming-sensitive neurons adapt to repetitive stimuli when ISIs are short. For ISIs of 1 or 3 s, neuronal responses exponentially declined with repetitive presentations, whereas responses remained stable at ISIs of 10 or 20 s (Fig. 2). Looming neuron adaptation has previously been done in locusts (Gray, 2005; Rind et al., 2008), zebrafish (Mancienne et al., 2021; Fotowat and Engert, 2023) and crabs (Oliva et al., 2007; Hemmi and Merkle, 2009). Interestingly, while crab lobula neurons adapt strongly when the ISI is 2 s (Berón De Astrada et al., 2013), i.e. on a similar time course to our data here, the locust DCMD adapts even when the ISI is very long, up to 80 s (Rind et al., 2008). In contrast, we saw no adaptation when the ISI was 10 or 20 s. This is interesting as the DCMD is also a descending neuron, and thus likely to have similar behavioral significance.

Similar time courses have been observed in zebrafish, where loom sensitive neural circuits adapt substantially when the ISI is 40 s (Fotowat and Engert, 2023). This is consistent with behavior, which adapts when the ISI is 10 s and 180 s (Fotowat and Engert, 2023). In insects, *Drosophila* escape jumps decrease substantially when an overhead looming stimulus is repeated 20 times over 5 min (Zacarias et al., 2018), whereas freezing behavior increases across repetitions (Zacarias et al., 2018). Thus, behavior appears to adapt even when the ISI is very long, inconsistent with our results (Fig. 5).

### Adaptation occurs primarily at the start of the response

We found that the neurons responded primarily before the time of maximum loom size (Fig. 4B). This is consistent with data from a range of looming neurons in insects (Gabbiani et al., 1999) and vertebrates (e.g. Fotowat and Engert, 2023). The onset of the response has been shown to be related to the angular size of the stimulus (Gabbiani et al., 1999, 2002; Fotowat and Gabbiani, 2011).

Our analysis of the temporal dynamics of neuronal responses revealed that adaptation primarily occurs at the onset of the response (Fig. 3). Indeed, the adaptation to the shorter ISIs (Fig. 2), appears to be driven by a shorter response duration (Fig. 3D, F), by primarily delaying the response onset (Fig. 3C, E). Adaptation taking place at the beginning of the response seems to be a conserved feature, as it was observed, but not quantified, before (Gray, 2005; Gray et al., 2010; Rosner and Homberg, 2013). In the future it would be interesting to use different *l/|v|* to investigate that the neurons shift their angular threshold with adaptation, or if there are other mechanisms at play. Nevertheless, an increased latency to the first spike has also been observed in blowfly H1 (Heitwerth et al., 2005), a neuron that responds to widefield optic flow.

### Adaptation occurs within the neuron itself

We found that adaptation likely takes place within the descending neuron itself, rather than in its pre-synaptic inputs (Fig. 4). Indeed, we reasoned that stimulation of pre-synaptic preferred-side inputs would be the same in unilateral and bilateral neurons (Fig. 4A), but that bilateral neurons would also be excited by non-preferred side input. As bilateral neurons were excited by stimuli on both sides, they would experience an effective ISI of 1 s (Fig. 4B, H), leading to strong adaptation (Fig. 4H, J, K). In contrast, as unilateral neurons only responded to the preferred side (Fig. 4F, H), they experienced a longer effective ISI (4 s), and no strong adaptation (Fig. 4H, I, K). Interestingly, locust DCMD neurons adapt at localized synapses (Gray 2005), and it was suggested that this allows the neurons to maintain sensitivity to stimuli outside their receptive field while adapting to repetitive inputs within it. Importantly, we show that bilateral neurons are still able to respond to looming stimuli on the non-preferred side (Fig. 4H). Indeed, as our experiments and subsequent analysis were performed differently, direct comparison is difficult.

Motion adaptation in lobula plate tangential cells (LPTCs) can be broken down into several components, a fast, local contrast gain reduction, a slow, global after-potential, and a slow, local output-range reduction (Harris et al., 2000; Nordström and O’Carroll, 2009; O’Carroll et al., 2011). Future work using different contrasts, different *l/|v|,* and different ISIs, can be used to investigate if looming adaptation in descending neurons can similarly be broken down into individual components, with individual time course and location.

### Behavioral responses to looming stimuli do not habituate

Despite the pronounced adaptation observed in looming-sensitive neurons (Fig. 1-3), the WBA of tethered hoverflies did not seem to adapt to repetitive looming stimuli. Instead, the hoverflies maintained robust WBA amplitudes across all ISIs tested (Fig. 5F). This discrepancy between neuronal (Fig. 1-3) and behavioral adaptation (Fig. 5) can be attributed to several factors. For example, when using the shorter ISIs (1 and 3 s), to match electrophysiological recordings, this interval was too short for the WBA to fully return to its baseline (green and orange data, Fig. 5F). However, when using longer ISIs, the WBA returned to baseline levels between repetitions (red and purple, Fig. 5F), but this ISI was too long to generate neuronal adaptation (see e.g. Fig. 2K).

Nevertheless, the apparent lack of behavioral adaptation (Fig. 5) highlights the complexity of sensorimotor integration. While individual neurons may adapt to repetitive stimuli, the overall behavioral output is shaped by the interplay of multiple neural circuits. Indeed, the WBA is affected by many descending neurons (Namiki et al., 2022), and not just the looming-sensitive neurons that we recorded from here. Therefore, other descending neurons may have controlled the WBA when the looming neurons had adapted. For instance, optic flow-sensitive descending neurons, which also respond to looming stimuli (Nicholas et al., 2020), could provide compensatory inputs to maintain consistent behavioral responses. Investigating their adaptation to looming stimuli could therefore be useful for understanding the integration of sensory inputs in the control of behavior. Nevertheless, the ability of multiple neurons to respond to looming stimuli could help ensuring robust responses to critical threats. Understanding how these circuits interact will be essential for unraveling the neural basis of adaptive behavior.

Interestingly, Rind et al., 2008 showed differences in locust DCMD adaptation to looming stimuli depending on behavioral state. When the locusts were flying rather than being physically passive, this greatly reduced DCMD adaptation. Therefore, the lack of behavioral adaptation that we observed here (Fig. 5) may reflect an underlying lack of neural adaptation, which we did not observe in electrophysiology as this was all done in immobilized animals. In any case, future research should explore the integration of inputs from different neuronal pathways and their contributions to behavioral responses.

## CRediT author statement

**Katja Sporar Klinge:** conceptualization, methodology, validation, formal analysis, investigation, data curation, writing – original draft, visualization; **Karin Nordström:** conceptualization, methodology, resources, writing – review & editing, supervision, project administration, funding acquisition.

## Acknowledgements

We thank Biomedical Engineering at SAHLN and the Botanic Gardens of Adelaide for their ongoing support. This research was funded by the US Air Force Office of Scientific Research (AFOSR, FA9550-23-1-0473) and the Australian Research Council (ARC, DP210100740 and DP230100006).

